# Behavioral Evidence for Enhanced Olfactory and Trigeminal Perception in Congenitally Deaf Individuals

**DOI:** 10.1101/2022.06.01.494382

**Authors:** Catherine Landry, Rim Nazar, Marie Simon, François Genest, Fanny Lécuyer Giguère, Johannes Frasnelli, Franco Lepore

**Affiliations:** Département de Psychologie, Université de Montréal, Montréal, QC, CAN; Research Institute of the MUHC, Montréal, QC, CAN; Département de Psychologie, Université de Sherbrooke, Sherbrooke, QC, CAN; Centre de recherche de l’hôpital Sacré-Coeur de Montréal, Montréal, QC, CAN; Département d’anatomie, Université du Québec à Trois-Rivières, Trois-Rivières, QC, CAN

**Author notes:** Contributed equally to the manuscript (Catherine Landry, Rim Nazar). **Author Note**.

**Keywords:** Deaf, Olfactory Perception, Trigeminal, Adaptation

## Abstract

Sensory deprivation, particularly hearing loss, is an excellent model to study neuroplasticity in the human brain and the adaptive behaviors that support the daily lives of deprived individuals. In adaptation to their hearing loss, deaf individuals rely on their other intact senses. Visual and tactile abilities are enhanced in deafness, but few studies have evaluated the olfactory function. This study aimed to compare the impact of congenital deafness on olfactory capacities using psychophysical tasks. Methodological issues raised in previous studies, such as homogeneous onset of deafness and cognitive function assessment, were considered. Eleven individuals with bilateral severe-to-profound deafness since birth were compared to 11 hearing non-signers similar in age (age range = 20-51 years old) and sex (7 women). The deaf subjects were assessed using various standardized neuropsychological tests to ascertain typical cognition. Olfactory functions were evaluated using the Sniffin’ Sticks battery test, which measures olfactory detection threshold, odor discrimination, and odor identification. Further, accuracy and response time were examined for the identification and localization of two odors to disentangle olfactory sensitivity from sensitivity in the trigeminal system. The Sniffin’ Sticks test demonstrated superior performances in the deaf participants to discriminate and identify odors. In line with this, they also showed higher sensitivity when both identifying and localizing odors. These findings suggest that congenital deafness is associated with superior performance in higher-level olfactory processing and increased sensitivity in the trigeminal system.

## Introduction

One approach to study neuroplasticity in the human brain and related adaptive behavior is through sensory deprivation. Adaptive behaviors support the daily life of individuals deprived of a sensory modality such as hearing using the other, intact senses (Alencar et al., 2019; Bavelier & Neville, 2002; Bell et al., 2019; Pavani & Bottari, 2012).Consequently, sensory loss plays a central role in the reorganization of functional processing in the intact sensory modalities, well known as neuroplasticity. Various animal (e.g., Lomber et al., 2010), psychophysical tasks (e.g., Megreya & Bindemann, 2017; Smittenaar et al., 2016), and brain imaging studies (e.g, Simon et al., 2020) demonstrate enhanced visual performances in the congenitally deaf. Although limited, the literature regarding the tactile modality presents superior sensitivity in relation to deafness when complex and cognitive tactile processes are studied (Sharp et al., 2020; van Dijik et al., 2013). Performance enhancement related to intermodal recruitment of the auditory and other sensory areas support the hypothesis of compensatory neuroplasticity. Active regions or pathways of the brain extend at the cost of less engaged regions or pathways (Rauschecker, 1995). Therefore, like these two modalities, it can be hypothesized that hearing loss also leads to enhancement of olfactory function. Hence, this study aims to evaluate if congenital deafness induces behavioral differences in olfactory and trigeminal abilities in comparison to hearing pairs.

To date, there is a general lack of studies that quantified chemosensory abilities of deaf individuals. Previous research showed reduced performance in profound hearing-impaired adults (Diekmann et al., 1994) and congenitally deaf adolescents (Guducu et al., 2016). However, a delay in language acquisition in deaf individuals might have influenced their understanding of the tasks, especially the congenitally deaf (Diekmann et al., 1994). In turn, congenitally deaf exhibit distinct cognitive abilities, such as a better capacity to direct visual attention during tasks (Colmenero et al., 2004; Parasnis & Samar, 1985) and better spatial memorization with deaf signers compared to hearing non-signers (Cattani & Clibbens, 2005). However, control for cognitive function was not included in earlier studies on olfactory ability in deaf individuals. This is crucial as olfactory tasks such as odor identification and discrimination are intrinsically associated with proficiency in executive functioning and semantic memory across the adult life span (e.g., Hedner et al. 2010; Larsson et al., 2000).

Another shortcoming in the literature is the predominance of olfactory dysfunction. Out of the thirteen deaf participants in one study (Guducu et al., 2016), three presented normosmia (normal olfaction), seven met hyposmia clinical criterion (reduced odor detection) and two participants ranked with anosmia (loss of smell). Although participants had taken an otorhinolaryngological examination to ensure the exclusion of sinonasal pathologies and nasal septal deviation, the prevalence of olfactory impairment is higher than expected in the population (Yang & Pinto, 2016), that is, 15% cases of hyposmia and 5% anosmia (Landis et al., 2009). What remains unclear is to what extent the poor scoring reported on psychophysical tasks relate to hearing loss, higher order cognitive deficiency/proficiency, inadequate sampling or, a possible lack of understanding the task instructions by the deaf sample (Guducu et al., 2016). In short, current knowledge regarding olfactory acuity in deafness is limited by significant methodological factors, such as heterogeneous clinical scoring and deafness onset, as well as lack of cognitive function measures. The duration of deafness and auditory rehabilitation by means of a cochlear implant (CI) are also factors known to influence the extent of neuroplasticity in impaired hearing individuals (Kral et al., 2016). Consequently, these factors were considered in the present research protocols.

Pure odorants that exclusively stimulate the olfactory system are rare. When evaluating the sense of smell, odorants most likely to be found in the environment are good experimental stimuli to use. However, most odorants have mixed olfactory-trigeminal components that can be detected even by individuals with anosmia (Doty et al., 1978). The trigeminal system processes sensations such as burning, cooling, itching, or stinging evoked my volatile substances (Laska et al., 1997). Only activation of the trigeminal nerve makes it possible to localize odors (Frasnelli et al., 2007; Frasnelli et al., 2009; Kleemann et al., 2009) i.e., determining which nostril is stimulated during monorhinal stimulations. In fact, chemical (as well as thermal) stimulation of the trigeminal nerve activates different polymodal ion channels from the transient receptor potential (TRP) subfamily (Frasnelli & Manescu, 2017; Viana, 2011). For example, cooling sensation induced by cool temperatures or chemical agents such as eucalyptol (eucalyptus) activate the TRPM8 receptor (Behrendt et al., 2004; Boonen et al., 2016).Other molecules stimulate the TRPA1 receptor, ranging from benzaldehyde (almond-like odor) (Richards et al., 2010) to various other irritants (Viana, 2011). In addition to its adaptative and protective functions for localizing potentially dangerous external stimuli, the trigeminal system plays an important role in the overall chemosensory experience. It may therefore be conceivable that trigeminal sensitivity is increased in congenitally deaf individuals. However, trigeminal sensitivity in deaf individuals has not yet been investigated.

This study explored olfactory and trigeminal processing behaviorally in congenitally deaf individuals. Specifically, detection threshold, discrimination and identification of odors was assessed, combined with a task of timed odor localization and identification. Addressing previous methodological limitations and controlling for cognitive function, olfactory performance is expected to be enhanced in individuals with congenital deafness given the crossmodally adaptative neuroplasticity in sensory deprivation. In line with visual and tactile performance, congenitally deaf individuals should manifest superior olfactory performances. This difference should extend to the trigeminal system, with deaf individuals presenting better ability to localize odors.

## Materials & Methods

### Participant Characteristics

Twenty-two participants took part in this study: 11 individuals [male = 4, female = 7; age range = 20-51 years (*M* = 35.64; *SD* = 9.63)] with severe-to-profound bilateral congenital hearing loss assessed by audiologists and 11 hearing non-signers [male = 4, female = 7; age range = 20-52 years (*M* = 35.64; *SD* = 10.42)]. The cause of deafness was sensorineural for all 11 deaf individuals. Experimental and control groups were matched for sex and age. Only individuals with a normal sense of smell were included (combined threshold-discrimination-identification (TDI) score ≥ 31/48). Individuals with psychiatric or neurological disorders (Moberg et al., 1999; Moscavitch et al., 2009), otolaryngology disease associated with olfactory dysfunctions (Landis et al., 2009), and smokers (Vennemann et al., 2008) were excluded. Every participant was instructed to avoid eating an hour before the experiment and were prohibited to use scented products on the day of testing. The study was approuved by the Research Ethics Board of the Centre de Recherche Interdisciplinaire en Réadaptation du Montréal métropolitain (CRIR) and the Centre de Recherche de l’Hôpital Sacré-Coeur de Montréal. All participants enrolled voluntarily and gave written informed consent. Participation was rewarded with monetary compensation. Table 1 presents the sociodemographic characteristics of all the participants.

**Table 1.** Descriptive Characteristics of Participants

| Id | Condition | Age<br>(years) | Gender | Education<br>(years) | Language<br>preference | CI duration<br>(years) |
| --- | --- | --- | --- | --- | --- | --- |
| 1 | D | 45 | F | 16 | LSQ + French | 42 |
| 2 | D | 28 | F | 14 | LSQ + French | 24 |
| 3 | D | 20 | M | 14 | LSQ + French | 18 |
| 4 | D | 28 | M | 16 | French | 27 |
| 5 | D | 47 | M | 9 | French | 42 |
| 6 | D | 51 | F | 7 | LSQ | N/A |
| 7 | D | 36 | F | 16 | LSQ | N/A |
| 8 | D | 32 | F | 14 | LSQ | N/A |
| 9 | D | 35 | M | 16 | LSQ | N/A |
| 10 | D | 28 | F | 16 | LSQ | N/A |
| 11 | D | 42 | F | 11 | LSQ | N/A |
| 12 | H | 46 | F | 11 | French | N/A |
| 13 | H | 47 | M | 16 | French | N/A |
| 14 | H | 52 | F | 13 | French | N/A |
| 15 | H | 36 | F | 16 | French | N/A |
| 16 | H | 31 | F | 16 | French | N/A |
| 17 | H | 20 | M | 14 | French | N/A |
| 18 | H | 34 | M | 14 | French | N/A |
| 19 | H | 28 | M | 16 | French | N/A |
| 20 | H | 27 | F | 21 | French | N/A |
| 21 | H | 26 | F | 18 | French | N/A |
| 22 | H | 45 | F | 12 | French | N/A |

*Note.* D = Deaf individuals; H = Hearing individuals; F = female; M = male; LSQ = Quebec Sign Language; CI = Cochlear implant; N/A = Not applicable.

### Cognitive Assessments

Given previous studies that suggest differential cognitive abilities in the deaf, cognitive function was ascertained in these participants by administering a series of validated non-verbal neuropsychological tests. Specifically, they included (1) spatial memorization capacities and visuo-constructive skills, specifically planning capacity, organizational skills, and perceptual and motor functions with the *Rey-Osterrieth Complex Figure test* (Deborah & Jane, 1985); (2) visuospatial and visuomotor coordination skills with the Blocks subtest and visuoperceptual and logical reasoning skills with the Matrix subtest of the *Wechsler Abbreviated Scale of Intelligence II* (Wechsler, 2013); and (3) ability to orient and maintain an adequate and stable level of efficiency throughout a visual activity with *Ruff 2 & 7 test* (Ruff et al., 1992). Each percentile rank score was within average when compared to the normative samples of the respective tests. More precisely, deaf individuals had average Z-scores for the Rey-Osterrieth Complex Figure test in immediate recall (Z = -0.67), delayed recall (Z = -0.6), and recognition (Z = 0.2), the Blocks test (Z = 1.09), the Matrix test (Z = 0.87) and the Ruff 2 & 7 test, both in accuracy (Z = -0.6) and speed (Z = 0.24) within the normative average.

### Procedure

Deaf participants were recruited through program managers, audiologists, and billboard advertisements at the Institut Raymond-Dewar of Montreal. The hearing participants were recruited online or through notices posted on the University of Montreal billboards. Control participants were chosen according to their age and sex to allow matching with deaf individuals. Information related to hearing loss, medical history and demographic information was collected through emails to ensure compliance with the inclusion criteria of the study. To address potential communication difficulties and promote optimal participation of deaf signers, a Quebec sign language (LSQ) interpreter translated the task instructions and the research protocol into LSQ. Videos of the interpreter were presented to the deaf participants before each tasks using a touchpad. The experimenter also communicated with the deaf participants through paper writing until complete understanding. Instructions were given verbally to the control participants.

#### Questionnaires

A short questionnaire was designed to ascertain the participants’ demographic information (education, age, sex). Deaf participants also filled two other questionnaires concerning their hearing impairment history (etiology, onset of deafness, duration of hearing loss, CI usage) and their means of communication (age exposure to language, oral fluency, and sign language).

#### Olfactory Function Assessment

Olfactory capacities were evaluated with the Sniffin’ Sticks test (SST; Burghardt, Wedel, Germany) (Hummel et al., 1997). This 40-to-60-minute validated task (Kobal et al., 2000; Hummel et al., 2007) uses a set of felt-tip pen-like odor dispensing devices – the Sniffin’ Sticks – designed to release odors at increasing intensity/different quality and evaluates threshold detection (T), discrimination (D) and identification (I) of odors. Three scores ranging from 1 to 16 were obtained for each condition. The TDI score (Wolfensberger, 2000) ranks the performance on a clinical scale from 1 to 48: normosmia (TDI ≥ 31), hyposmia (15 < TDI < 31) and anosmia (TDI ≤ 15) (Hummel et al. 2007; updated version Oleszkieicz et al., 2019). Participants were blindfolded to avoid visual identification of odors through stimulus recognition. After each trial, participants were asked to validate the response written on paper. The following sections briefly detail the assessment procedure of each subtask (for more details, see Rumeau & Jankowski, 2016).

1-The *detection* threshold (T) was assessed with a 3 multiple forced choice following an ascending/descending staircase procedure to 16 triplets of sticks. For each trial, three sticks were presented (20 seconds each) in random order, two containing a solvent and the third containing a diluted concentration of phenylethanol alcohol (PEA, rose-like odor) according to predefined degrees. Participants were asked to identify the PEA stick within the triplets. Two consecutive correct identifications of the PEA stick reverse the staircase to a lower concentration staircase. An error reverses the scale to a higher concentration staircase. The test ends when the scale reversal criterion is encountered seven times. The detection threshold score was defined as the mean of the last four staircase turns (Hummel et al., 2007).

2-The *discrimination* task (D) uses 16 triplets of sticks randomly presented, two sticks containing the same odorant and a third one containing the target odorant (Hummel et al. 2007). Participants were asked to determine which of the three sticks smelled differently.Each triplet was separated by at least 30 s and each stick presentation was separated by a 3 s interval. The discrimination score results in the sum of correctly identified odd sticks.

3-The odor *identification* (I) was carried out using a 4 multiple-forced choice where the participant had to correctly identify 16 sticks containing different common odors (Hummel et al. 2007). Participants could freely smell as much as considered necessary before answering. Each stick presentation was separated by a 30 s interval. The identification score is defined by the sum of correctly identified odors.

#### Automated Odorant Localization and Identification (AOLI)

A computer-controlled device delivering fast and stable stimulus (Lundström et al., 2010) was used to measure the identification and localization of odors. A total of 36 stimulations were presented: 12 air (control), 12 benzaldehyde [almond (A); Sigma– Aldrich, St. Louis, MO, USA] and 12 eucalyptol [eucalyptus (E); Galenova, St.- Hyacinth, QC, Canada] stimuli. Benzaldehyde and eucalyptol were chosen because they are mixed olfactory-trigeminal stimuli, i.e., they stimulate both the olfactory and the trigeminal system. Both benzaldehyde and eucalyptol were diluted at 50% in propylene glycol (Sigma–Aldrich, St. Louis, MO, USA). Stimulations were delivered in a randomized order through one of the two nostrils: [left (L) or right (R)] every 30 s. On half of the trials, the participant was asked the command “Where?” (localization of the stimulated nostril, R or L; forced choice); the other half they were asked the command “What?” (identification of the delivered stimuli, A or E; forced choice) before odorant delivery via computer screen (Kéïta et al., 2013). By doing so, sensitivity in the trigeminal system (odor localization) and in the olfactory system (odor identification) were assessed. The participants were instructed to press the adequate keyboard button as fast as possible. The number of hits (correct detection when the stimulus is present), false alarms (inaccurate detection of the stimulus when absent) and response time (in seconds) was measured. To avoid habituation, stimuli were separated by a 40 s fixed interval and conditions were counterbalanced. A white noise was continuously played to cover the sounds emitted by the device to prevent cueing the localization or identification of odorants. This preventive procedure was perpetuated in the deaf group for protocol standardization purposes.

The sensitivity index *d’* and response bias *C* were then computed based on the number of hits and false alarms (Stanislaw & Todorov, 1999). In Signal Detection Theory, the sensitivity index represents the accuracy with which the stimulus can be detected by comparing hits with false alarms. The higher the values of *d’*, the better the task was performed. The response bias *C* indicates the participant’s subjective criterion for responding: a positive bias reveals a tendency either towards a side (localization task) or stimulus (identification task).

### Statistical Analyses

Data was analyzed with SPSS (version 27 for macOS 10.14+, SPSS Inc., Chicago, IL). Sphericity (Mauchly’s test of Sphericity, *p* > 0.05), normality (Kolmogorov-Smirnov test for normality, *p* > 0.05) and variance homogeneity (Levene’s test, *p* > 0.05) postulates can be assumed for SST and AOLI scores data.

An independent sample *t*-test was performed to compare the computed TDI score between deaf and hearing individuals. A repeated-measures analysis of variance was next carried out (rm-ANOVA) to compare the effect of *condition* (between-subject factor; 2 levels: deaf, hearing) on scores with *task* (3 levels: detection, discrimination, identification) as within-subject factor. For the AOLI scores, the dependent variables sensitivity index and response bias were submitted to a rm-ANOVA with *task* (2 levels: identification, localization) as within-subjects factor and *condition* (2 levels: deaf, control) as between subjects factor.

Finally, mean scores (SST, AOLI) were subjected to the Mann-Whitney *U* test to compare deaf individuals wearing a CI at the time of testing with those who have no CI experience. All post-hoc comparisons were corrected with Bonferroni’s procedure. The alpha level of significance was set at 0.05 for all analyses.

## Results

### Olfactory Function Assessment

On average, deaf participants (*M* = 40.159, *SD* = 3.204) obtained a higher global TDI score than control individuals (*M* = 36.204, *SD* = 3.409). This difference (95% CI, 0.270-2.096) was significant, *t*(20) = -2.803, *p* = 0.011 with a high effect size, *d* = 1.195.

With regards to the individual tests of the SST, the rm-ANOVA revealed no statistically significant interaction between *task* * *condition* (*F*(2,40) = 0.463, *p* = 0.633, *η*^*2*^ = 0.23). A significant main effect of *condition* was observed (*F*(1,20) = 7,859, *p* = 0.011, *η* ^*2*^ = 0.282), with deaf individuals outperforming controls, in line with the result of the global TDI score. Finally, we observed a significant main effect of *task* (*F*(2,40) = 9.795, *p* < 0.001, *η*^*2*^ = 0.329), with highest scores being reached for the identification task, followed by the discrimination task, and the olfactory threshold task (Figure 1, Table 2).

**Figure 1.**
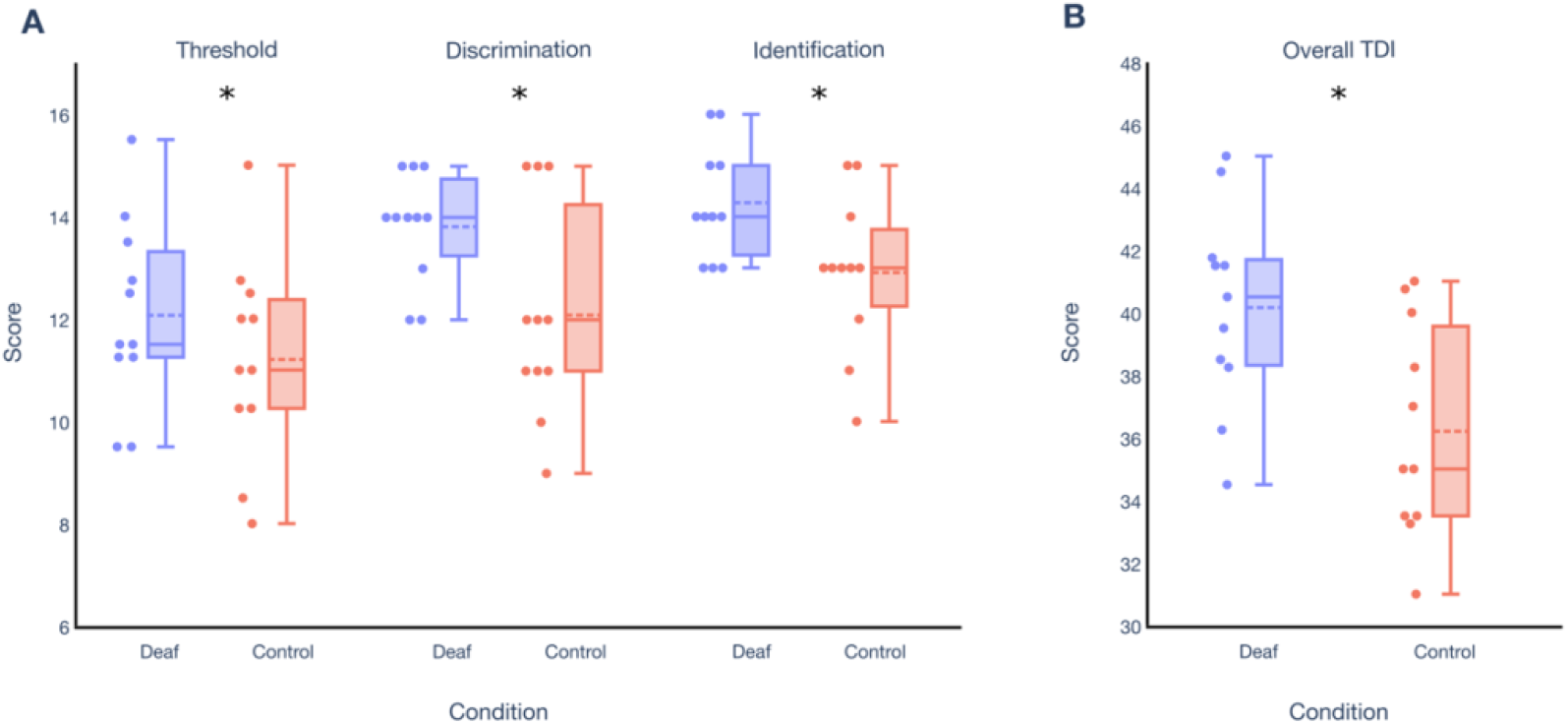
Mean scores Distribution on the Sniffin’ Sticks Test, Separated by Condition (Deaf, Control) and Tasks (Olfactory Threshold, Discrimination, Identification) and Overall TDI *Note*. ^A^Olfactory detection threshold, discrimination, and identification scores, each out of 16 and ^B^Overall TDI, score out of 48. Asterisk indicates a significant group effect at *p* < 0.05. The dashed line in the box represents the mean value, while the solid line represents the median. The first and third quartile are delimited by the lower and upper horizontal line of the boxplot. The minimum and maximum values are indicated by the lower and upper fences, respectively. Each point designates the performance of one participant.

**Table 2.** Sniffin’ Sticks Test Descriptive Statistics Separated by Task (Threshold, Discrimination, Identification) and Condition (Deaf, Control)

|  | Threshold |  | Discrimination |  | Identification |  |
| --- | --- | --- | --- | --- | --- | --- |
|  | Deaf | Control | Deaf | Control | Deaf | Control |
| *N* | 11 | 11 | 11 | 11 | 11 | 11 |
| *Mean* | 12.068 | 11.205 | 13.818 | 12.091 | 14.273 | 12.909 |
| *SD* | 1.827 | 1.981 | 1.079 | 2.071 | 1.104 | 1.514 |
| *SE* | 0.575 | 0.575 | 0.498 | 0.498 | 0.399 | 0.399 |
| *Lower limit* | 10.87 | 10.006 | 12.780 | 11.052 | 13.440 | 12.076 |
| *Upper limit* | 13.267 | 12.403 | 14.857 | 13.130 | 15.106 | 13.742 |
| *P* | 0.011 |  |  |  |  |  |

### Automated Odorant Identification and Localization (AOLI)

For the sensitivity index *d’*, the rm-ANOVA yielded no significant interaction between *task * condition* (*F*(1,20) = 0.009, *p* = 0.924, *η*^*2*^ = 0.00). There was a large main effect of *task* (*F*(1,20) = 10,71, *p* = 0.004, *η*^*2*^ = 0.349). On average, participants performed more accurately in the identification (*M* = 1.057, *SD* = 0.666) compared to localization (*M* = 0.361, *SD* = 0.856). A significant main effect of *condition* was also observed (*F*(1,20) = 5.652, *p* = 0.028, *η*^*2*^ = 0.22). Deaf individuals outperformed hearing controls in both tasks, i.e., identification and localization (Table 3).

**Table 3.** Automated Identification and Localization Descriptive Statistics Separated by Measure (Sensitivity Index, Response Bias) and Condition (Deaf, Control)

|  | Identification |  |  |  | Localization |  |  |  |
| --- | --- | --- | --- | --- | --- | --- | --- | --- |
|  | Sensitivity index |  | Response bias |  | Sensitivity index |  | Response bias |  |
|  | Deaf | Control | Deaf | Control | Deaf | Control | Deaf | Control |
| *N* | 11 | 11 | 11 | 11 | 11 | 11 | 11 | 11 |
| *Mean* | 1.339 | 0.775 | 0.227 | 0.280 | 0.622 | 0.100 | 0.178 | 0.309 |
| *SD* | 0.277 | 0.824 | 0.257 | 0.350 | 0.827 | 0.838 | 0.603 | 0.546 |
| *SE* | 0.185 | 0.185 | 0.093 | 0.093 | 0.251 | 0.251 | 0.173 | 0.173 |
| *Lower limit* | 0.952 | 0.388 | 0.034 | 0.087 | 0.98 | -0.424 | -0.183 | -0.053 |
| *Upper limit* | 1.725 | 1.162 | 0.420 | 0.473 | 1.146 | 0.623 | 0.540 | 0.670 |
| *P* | 0.028 |  |  |  |  |  |  |  |

On average, response bias C was slightly positive for the identification (*M* = 0.253, *SD* = 0.301) and localization (*M* = 0.243, *SD* = 0.565). No significant main effect was observed for *task* (*F*(1,20) = 0.005, *p* = 0.943, *η*^*2*^ = 0.00) or *condition* (*F*(1,20) = 0.421, *p* = 0.524, *η*^*2*^ = 0.021) or interaction *task * condition* (*F*(1,20) = 0.079, *p* = 0.782, *η*^*2*^ = 0.004).

On average, response time was similar for localization (*M* = 2.124, *SD* = 0.627) and identification (*M* = 2.056, *SD* = 0.686). Again, no effect of *task* (*F*(1,20) = 3.14, *p* = 0.083, *η*^*2*^ = 0.015) or interaction *task * condition* (*F*(1,20) = 0.32, *p* = 0.860, *η*^*2*^ = 0.002) were found. However, there was a main effect of *condition* (*F*(1,20) = 6,001, *p* = 0.024, *η*^*2*^ = 0.231), with deaf participants (*M* = 1.765, *SD* = 0.328) being faster at accurately identifying odorants compared to hearing controls (*M* = 2.346, *SD* = 0.833). This tendency extends to the localization task: deaf individuals (*M* = 1.856, *SD* = 0.436) were significantly faster in response time than control participants (*M* = 2.392, *SD* = 0.691). Results are illustrated in Figure 2.

**Figure 2.**
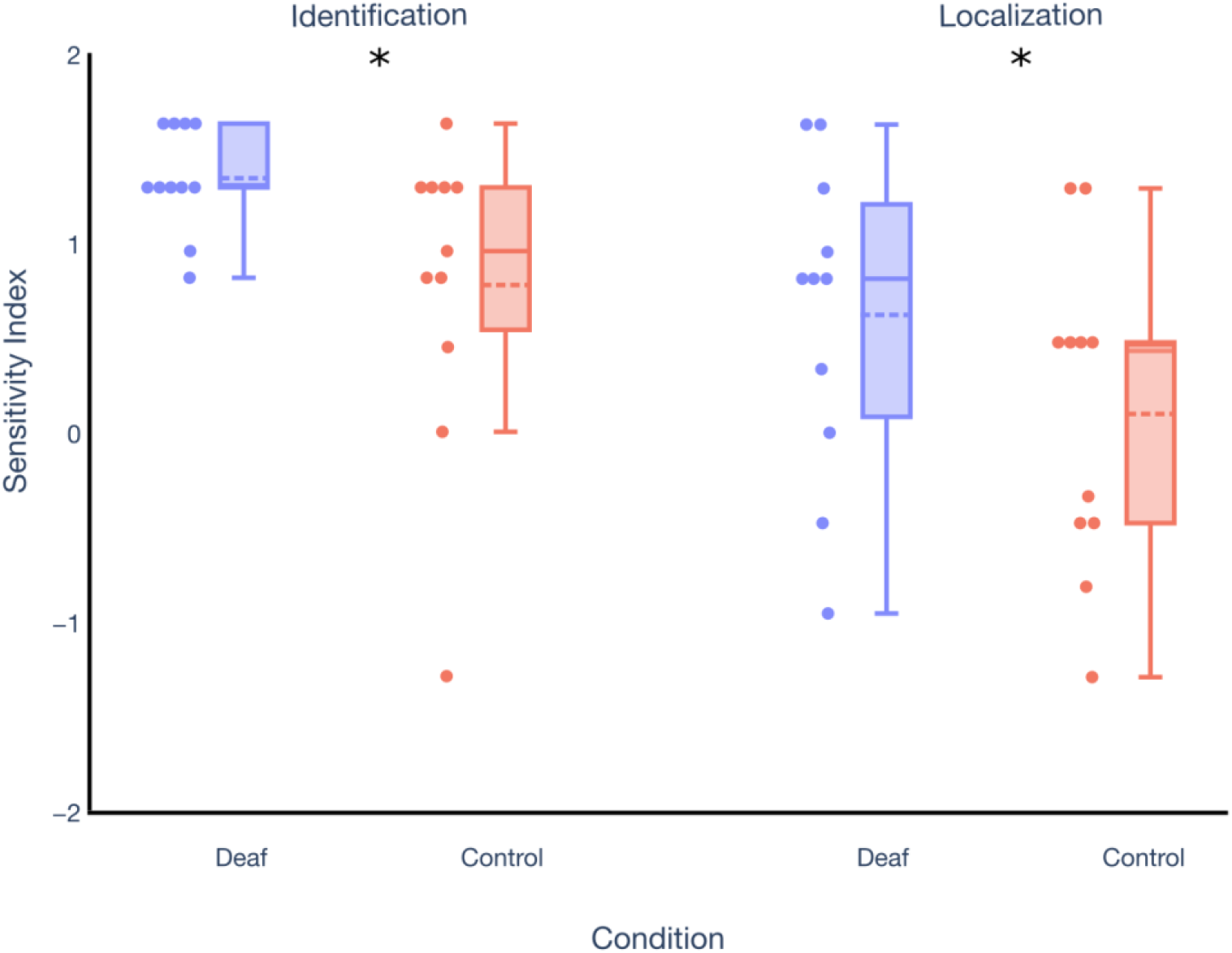
Sensitivity Index d’ Separated by Condition (Deaf, Control) and Task (Odor Identification, Odor Localization) *Note*. Asterisk indicates a significant group effect at *p* < 0.05. The dashed line in the box represents the mean value, while the solid line represents the median. The first and third quartile are delimited by the lower and upper horizontal line of the boxplot. The minimum and maximum values are indicated by the lower and upper fences, respectively. Each point designates the performance of one participant.

### Effect of Cochlear Implant

No significant difference was observed between CI users and those without CI for the Sniffin’ Sticks test global score (*U* = 13.5, *p* = 0.792), AOLI identification (*d’*: *U* = 9.5, *p* = 0.329; *C*: *U* = 25, *p* = 0.082) localization (*d’*: *U* = 20.5, *p* = 0.329; *C*: *U* = 18, *p* = 0.662) or response times: identification (*U* = 7, *p* = 0.117) or localization (*U* = 14, *p* = 0.931).

## Discussion

The present study examined the impact of congenital hearing loss on several chemosensory tasks. As hypothesized, olfactory abilities and trigeminal perception are enhanced for deaf individuals compared to hearing controls in discrimination and identification tasks.

This is in line with the notion of adaptive neuroplasticity between modalities, as observed for the visual and tactile domains in previous studies (Megreya & Bindemann, 2017; Sharp et al., 2020; Simon et al., 2020; Smittenaar et al., 2016; van Dijik et al., 2013). The results are however also in contrast with some existing literature on the olfactory modality showing reduced performance for the deaf. One specific aspect may explain this difference: in contrast to earlier reports (Guducu et al., 2016; Diekmann et al., 1994), we ascertained typical cognitive function in the deaf group. In fact, only deaf individuals were included who had scores within the normative range for attention, visual memory, and fluid reasoning. This is especially important because the ability to identify and discriminate odors correlates with performance in executive function and semantic memory (e.g., Hedner et al. 2010; Larsson et al., 2000). Either altered cognitive function or a delayed language acquisition might in fact influence previous results and explain some variability (Diekmann et al., 1994), mainly for the preponderance of hyposmia and anosmia in the deaf population (Guducu et al., 2016). In fact, vocabulary knowledge is particularly important in the context of the SST identification task, as it implies a forced choice labeling of odorants (Hummel et al., 1997). Future research with individuals having limited verbal language and reading abilities therefore should use appropriate tests and adapt the means used to present the response choices (written vs signed) to ensure full comprehension of the task.

The differences with the existing literature may also partly be explained by the age of onset deafness, a factor known to influence compensatory plasticity (Lazzouni & Lepore, 2014). There is a critical period at the start of postnatal life during which the brain is most receptive to change (Oberman & Pascual-Leone, 2013). However, heterogeneity of background within the deaf sample are common limitations in the literature (Bavelier et al., 2006). In one study (Diekmann et al., 1994), only three individuals were congenitally deaf while four became profoundly deaf during the first five years. In contrast, in the present study, only participants with congenital deafness, i.e., hearing loss present at birth, were included.

Our findings therefore provide support for the notion that crossmodal sensory compensation for the auditory system extends to chemosensory domains, by showing that the sense of smell is enhanced in deaf individuals. In agreement with this hypothesis, additional contribution of neural resources to the remaining sensory streams compensate for auditory deprivation (Singh et al., 2018). Cross-modal reorganization underlies adaptive and compensatory behaviors (Merabet & Pascual-Leone, 2010). While it is accepted that sensory deprivation leads to the expansion of other senses, these compensatory effects remain complex (Kupers & Ptito, 2014). For the olfactory modality, the neurobiological underpinnings are yet to be elucidated. Future studies using functional magnetic resonance imaging (fMRI) will possibly identify the implicated brain areas associated with improved olfactory performance as the result of compensatory neuroplasticity between auditory and olfactory modalities in deafness.

The superiority of deaf individuals includes the trigeminal system, as they had superior abilities to localize odorants. In fact, the trigeminal system is a third chemosensory system next to smell and taste. The trigeminal system has a protective function, which is reflected in various physiological reflexes such as salivation, tearing, coughing, respiratory depression, and sneezing (Viana, 2011). Its sensitivity can be assessed with the odor localization task used in the present study and validated by past ones (Frasnelli et al., 2007). This observation is also in line with results on individuals with early blindness (Manescu et al., 2021).

In addition to better response sensitivity, the results of this study also indicate that deaf individuals responded faster on the localization and identification tasks, the latter correlating with response accuracy (Kéïta et al., 2013). There is evidence that sensory deprivation affects the visual function, especially spatial information presented at the periphery, resulting in faster reaction time to stimuli (Bottari et al., 2011; Chen et al., 2006; Codina et al., 2017; Nava et al., 2008; Prasad et al., 2017). This advantage appears to emerge in adolescence and transfer to adulthood (Codina et al., 2011). It is possible that response time in olfactory function constitutes a behavioral compensation for deafness-induced neuroplasticity, similar to what is found in the visual modality.

One possible shortcoming of our study is the fact that five of the eleven deaf individuals were implanted with a CIs at the time of testing. Three of them were implanted before the age of 3.5, meaning that they had access to some form of auditory input during the sensitive period when the central auditory system is most receptive to change (Sharma et al., 2002). To ascertain that this was not a confounding factor, the five CI users were compared with the rest of the deaf sample and no statistically significant differences were found. However, in future studies, CI users will be treated as a separate group, as there are implications for the means of communication (oral or signed) and the degree of neuroplasticity. Larger sample sizes would allow to draw stronger conclusions about the use of CI and age of implantation on olfactory/trigeminal functions, as much as to confirm the present findings. It is to be noted that the handedness of participants is only available for controls, all of whom were right-handed.

## Conclusion

Psychophysical olfactory assessment methods were used to evaluate the influence of congenital hearing loss on chemosensory systems. Compared to controls, individuals with severe-to-profound congenital deafness but typical cognition had significantly higher olfactory scores. In addition, they were more sensitive to trigeminal odor localization.

## Acknowledgments

This research was supported by the Canada Research Chair Program and the Natural Sciences and Engineering Research Council of Canada grants (#RGPIN- 8245-2014, F.L.) and the Canadian Institutes of Health Research (#166197, F.L.).

